# Structural insights into the recognition of RALF peptides by FERONIA receptor kinase during Brassicaceae Pollination

**DOI:** 10.1101/2024.07.02.601642

**Authors:** Hemal Bhalla, Karthik Sudarsanam, Ashutosh Srivastava, Subramanian Sankaranarayanan

## Abstract

Ensuring species integrity and successful reproduction is pivotal for the survival of angiosperms. Members of Brassicaceae family employ a “lock and key” mechanism involving stigmatic (sRALFs) and pollen RALFs (pRALFs) binding to FERONIA, a *Catharanthus roseus* receptor-like kinase 1-like (CrRLK1L) receptor, to establish a prezygotic hybridization barrier. In the absence of compatible pRALFs, sRALFs bind to FERONIA, inducing a lock state for pollen tube penetration. Conversely, compatible pRALFs act as a key, facilitating successful fertilization. Competing pRALFs reduce the sRALFs binding to FERONIA in a dose-dependent manner, enabling pollen tube penetration. Despite its crucial role in Brassicaceae hybridization, the structural basis of this binding remains elusive owing to the highly flexible nature of RALF peptides. Using advanced structural modeling techniques and flexible peptide molecular docking, this study reveals that pRALFs and sRALFs bind to negatively charged pockets in FERONIA with varying binding affinities. Our study unveils the structural basis of this binding, shedding light on the molecular mechanism underlying hybridization barriers in Brassicaceae.

## Introduction

Flowering plants employ diverse reproductive barriers to maintain species integrity across interspecific and intergeneric hybridization. These barriers ensure successful fertilization, viable offspring production, and genetic isolation between species. Within Brassicaceae, a pre-zygotic hybridization barrier is established through a sophisticated “lock and key” mechanism at the pollen-pistil interface. This system involves the interaction of autocrine secreted stigmatic sRALFs (Rapid Alkalinization Factors) with the stigma membrane-associated CrRLK1L (*Catharanthus roseus* receptor-like kinase 1-like) receptor, FERONIA, which acts as a lock preventing pollen tube penetration and fertilization. Conversely, pollen-borne pRALFs from the same species serve as keys, facilitating successful pollen tube penetration and fertilization. (Figure 1A) (Lan et al., 2023). The binding between RALFs (stigmatic and pollen) with FERONIA is highly dynamic and dose-dependent, requiring higher doses of pRALFs to overcome the hybridization barrier established by sRALFs and FERONIA (Lan et al., 2023).

**Figure 1.**
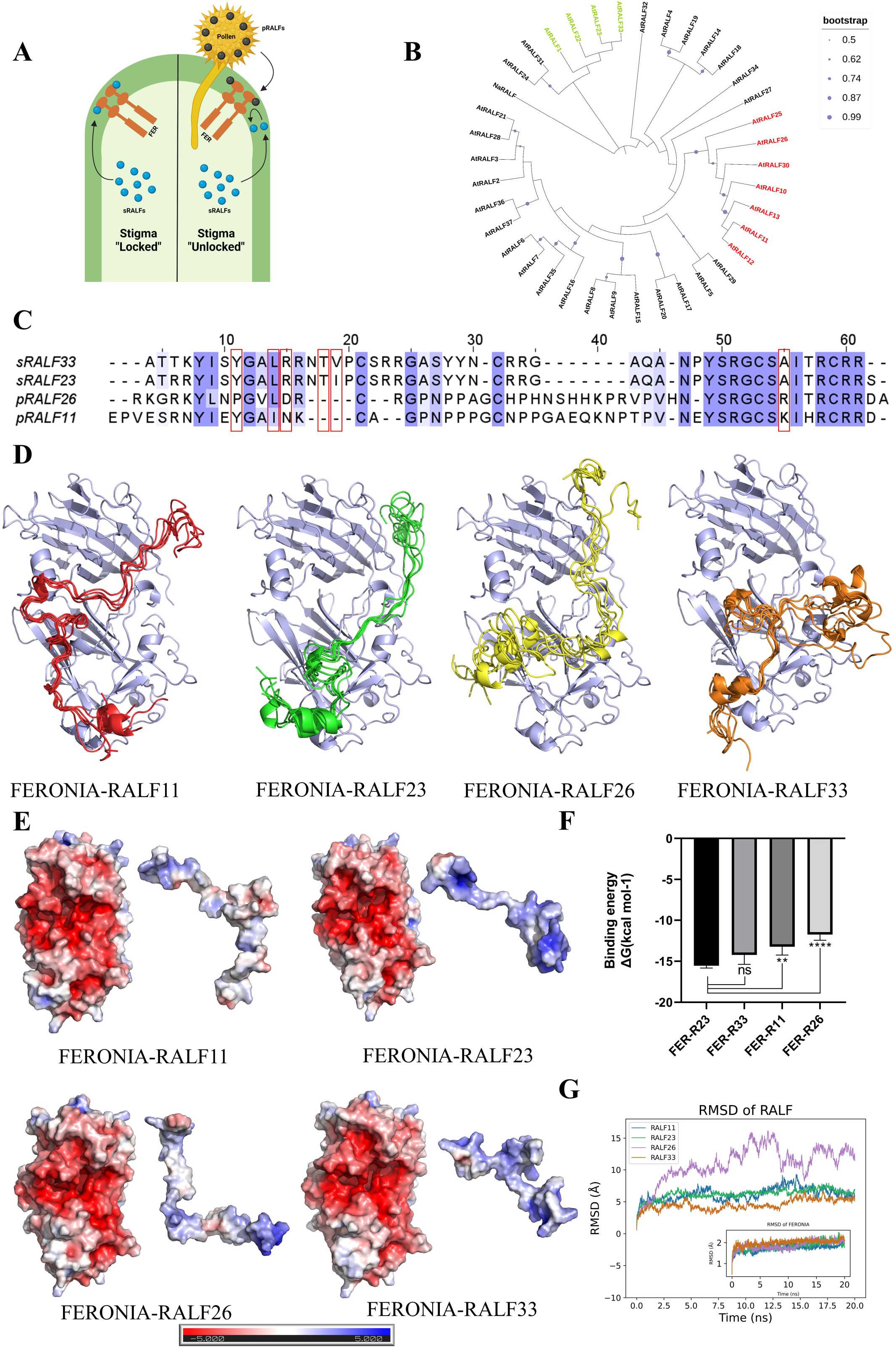
Structural insights into binding of various stigmatic and pollen RALFs to FERONIA. (A) Schematic diagram showing binding of stigmatic (sRALFs) and pollen (pRALFs) RALFs with the stigmatic membrane-associated receptor-like kinase FERONIA (FER). (B) Maximum likelihood tree inferred from multiple sequence alignment of AtRALF peptide sequences and *Nicotiana attenuata* RALF (NaRALF). (Supplementary Figure S1A). The tree was rooted using NaRALF. AtRALFs highlighted in green represent stigmatic RALFs, and red represent pollen RALFs. Bootstrap values (1000 repetitions) above 50% are represented by purple circles in the corresponding branches. (C) Alignment of sRALF23/33 and pRALF11/26 color-coded based on sequence conservation. The darker the color, the more conserved the residue. The Red box indicates residues used for unambiguous restraints during docking with FERONIA. (D) Superimposed top 5 structures of FERONIA-RALF complexes from the largest cluster predicted using FlexpepDock *ab initio* protocol and clustered using Calibur. (Blue: FERONIA, Red: RALF11, Green: RALF23, Yellow: RALF26, Orange: RALF33) (E) Representative structures of FERONIA-RALF models from the largest cluster, colored based on electrostatic potential as calculated by APBS. Each complex shows electrostatic potential at the interacting surface of FERONIA (left) and RALF (right). Red color indicates negative potential, and blue indicates positive potential. The darker the color, the more electrostatic potential. (F) Graph representing the binding energies of top 5 scoring structures of FERONIA-RALFs from each largest cluster predicted using Prodigy server. (Error bar indicates standard deviation, One way ANOVA, *P<0.05; n=5) (G) Root mean square deviation (RMSD) of C-α atoms of RALFs in RALF-FERONIA complexes across 20 ns MD simulation trajectory. The inset figure shows the RMSD of C-α atoms of FERONIA along the trajectory.

Various RALFs, including AtRALF1 and AtRALF23, are known to interact with the ectodomains of CrRLK1L receptors such as FERONIA (Haruta et al., 2014; Liu et al., 2018; Xiao et al., 2019; Liu et al., 2021). Experimental truncation studies have confirmed conserved motifs at both the N and C terminals of RALF peptides to be critical for their functional activity (Pearce et al., 2010). These peptides are known to adopt structured forms and form disulfide bonds across conserved cysteine residues either during or immediately preceding receptor binding (Pearce et al., 2010; Xiao et al., 2019).

The stigmatic RALF, AtRALF23, has been crystallized with FERONIA and LORELEI-LIKE GLYCOSYLPHOSPHATIDYLINOSITOL-ANCHORED PROTEIN 1 (LLG2), although the crystal structure resolved only the N-terminal region of AtRALF23 due to the flexibility of the C-terminal portion (Xiao et al., 2019). Structural insights into the binding of these RALFs with FERONIA remain limited due to unresolved experimental structures and the inherent flexibility of the peptides involved (Liu et al., 2018; Frederick et al., 2019; Xiao et al., 2019). Understanding these structures is crucial for elucidating the interactions between RALFs (stigmatic and pollen) with FERONIA, as well as for discerning the structural basis of recognition mechanisms that could potentially control interspecific and intergeneric hybridization. Furthermore, structural modeling and prediction of these complexes have been challenging due to the lack of structural restraints, the peptides’ flexible nature, and their unique propensity to adopt structure only upon or during complex formation with their respective receptors. In this study, structural modeling and flexible peptide docking have been utilized to gain insights into the binding of FERONIA-RALF complexes involved in maintaining the hybridization barrier in Brassicaceae.

## Results and Discussion

To investigate the interaction between RALFs (stigmatic and pollen) and FERONIA, we modeled the structures of these complexes using the Rosetta flexible peptide docking *ab initio* protocol (Raveh et al., 2011). In order to understand the phylogenetic relationships across various RALFs in *Arabidopsis thaliana*, sequences were retrieved from NCBI (see Table S1) (Sayers et al., 2021). Following multiple sequence alignment (MSA) (Supplementary Figure 1A and 1B), a phylogenetic tree was constructed (Figure 1B). MSA and phylogenetic analysis revealed known conserved motifs across various AtRALFs as well as motifs that are specifically present in sRALFs, such as YISY and ASYYN motifs (Lan et al., 2023). We also observed conserved motifs in pRALFs, such as PNPPPG and PVNE (Supplementary Figure 1B). Notably, sRALFs and pRALFs formed two separate clades in the AtRALF phylogeny in accordance with the specific conserved motifs seen in MSA, highlighting their functional significance. As seen in Figure 1B amongst stigmatic RALFs, RALF1 clusters with RALF23 and RALF33. Interestingly, RALF1, a close homolog of RALF23/33, is known to interact with FERONIA in the roots (Haruta et al., 2014; Liu et al., 2018). Although an experimentally derived structure is available for the FERONIA-RALF23 (PDB ID 6A5E) complex, the C-terminal region of RALF23 was unresolved due to its high flexibility in the crystal (Xiao et al., 2019). Bis(sulfosuccinimidyl)suberate (BS3) crosslinking and 1-ethyl-3-[3-(dimethylamino) propyl] carbodiimide hydrochloride (EDC)/glycine ethyl ester (GEE) labeling and mass spectrometry had previously revealed the interaction interface between RALF1 and FERONIA (Liu et al., 2018).

Secondary structure prediction indicated high-confidence coils for sRALF23/33 and pRALF11/26 (Supplementary Figure 2), similar to the previously known solution structure of RALF8, which showed highly unstructured peptide except in the vicinity of disulfide bridges (Frederick et al., 2019). This characteristic can be attributed to proline-rich regions, specifically in pollen RALFs, as well as the presence of cysteine residues within very short intervals leading to minimal peptide loop length between the disulfide bridge, which is known to play a key role in structural conformation. The tertiary structures predicted by AlphaFold (Supplementary Figure 3) exhibited consistent similarity to secondary structure predicted data showing extensive coiled regions in both stigmatic and pollen RALF structures. Attributing to the presence of coils with high confidence, the pLDDT (predicted local distance difference test) scores exceeded 60%, and PAE (predicted aligned error) scores were very low across all predicted models. Initial molecular docking simulations using Haddock 2.4 utilized the experimentally solved FERONIA structure, incorporating ambiguous and unambiguous restraints from the FERONIA-RALF23 crystal structure (Xiao et al., 2019) and interaction data from FERONIA-RALF1 (see Supplementary Table S2) (Figure 1C) (Liu et al., 2018). Clusters generated by Haddock 2.4 were evaluated based on cluster size, Haddock score, and root mean square deviation (RMSD). The selected FERONIA-RALF models predominantly belonged to cluster 1, except for FERONIA-RALF26, where cluster 3 was preferred due to its higher score and comparative RMSD to cluster 1 (see Supplementary Table S3, Supplementary Figure 4). These initial models were used for flexible peptide docking guided by consistent restraints (see Supplementary Table S2), wherein simultaneous folding of RALF peptide along with docking was performed in accordance with the notion that these peptides tend to fold upon binding to their receptors (Figure 1D and Supplementary Figure 5C) (Pearce et al., 2010; Xiao et al., 2019). The predicted models had an average score ranging from -295.24 to -534.182 and RMSD from 1.212 to 2.384 for the top 5 models out of 50,000 predictions (see Supplementary Table S4). Scores for the top 5,000 models were between -200 and -500, with RMSD values confined between 0 and 10, indicating significant scoring and RMSD uniformity across the predictions (Supplementary Figure 5A). Cluster analysis of these predicted models clearly indicated that cluster 1 predominated over cluster 2 in terms of representation, thus warranting its selection for further analysis (see Supplementary Table S5). Scores and RMSD values within cluster 1 of the FERONIA-RALF models demonstrated significant consistency, supporting their suitability for detailed examination (Supplementary Figure 5B). Notably, electrostatic calculations using APBS (Adaptive Poisson-Boltzmann Solver) revealed that RALFs, characterized by numerous positively charged residues, bind to a highly negatively charged pocket in FERONIA (Figure 1E and Supplementary Figure 5D). Binding energy analysis revealed that FERONIA-RALF23 and RALF33 (stigmatic RALFs) have similar binding energies, whereas FERONIA-RALF11 and RALF26 (pollen RALFs) have significantly lower binding energies, suggesting weaker binding to FERONIA (Figure 1F) (see Supplementary Table S6). Interaction studies revealed several polar contacts (see Supplementary Table S7). Interestingly, FERONIA, with aspartic acid and asparagine residues at positions 222 and 206, respectively, formed polar contacts with arginine at position 30 in both the stigmatic RALFs, i.e., sRALF23/33, whereas they were replaced by proline in pRALF11/26 (Supplementary Figure 6A and 6B). The lower binding energies of pRALFs with FERONIA can be attributed to their reduced number of positively charged residues and an increased number of negatively charged residues compared to sRALFs (Supplementary Figure 6C). Additionally, the highly coiled-coil structures of pRALFs, rich in proline which acts as a helix breaker, contrast with the structures of sRALFs (Supplementary Figure 6D). The presence of proline residues at sites of polar contacts can significantly influence the binding affinity to FERONIA. Molecular dynamics simulations further support our predicted models and binding energy analysis, demonstrating higher stability of FERONIA-RALF23/33 (sRALFs) compared to FERONIA-RALF11/26 (pRALFs) (Figure 1G and Supplementary Figure 7A). Specifically, pRALF26 exhibits the lowest stability (Figure 1G) and a highly unstable C-terminal domain (Supplementary Figure 7A), consistent with the binding energy data (Figure 1F). FERONIA structure remains stable throughout the trajectories in all four systems (Figure 1G, inset). During MD simulations, the distance between the C-α atoms of FERONIA D222 and R30, as well as N206 and R30 (discussed above), remained stable around 6.5 Å and 7 Å in the case of FERONIA-sRALF33 complex. For FERONIA-sRALF23 complex simulations, the two distances remained stable for the initial part of the simulation but increased later, suggesting a relatively weaker interaction (Supplementary Figure 7B and 7C). This observation supports the hypothesis that these residues may play a crucial role in the binding of stigmatic RALFs with FERONIA.

We have successfully predicted previously uncharacterized structural models for FERONIA and RALFs (stigmatic RALF23/33 and pollen RALF11/26), which are crucial for understanding the molecular details of the “lock and key” mechanism that serves as a hybridization barrier. This was achieved through a hybrid approach integrating experimental restraints into modeling highly flexible peptide-protein interactions (Supplementary Figure 8). Pollen tube penetration assays and *in vitro* co-immunoprecipitation assays have previously shown dose-dependent displacement of sRALF from FERONIA by considerably higher (100 times) doses of pRALF, suggesting differential interaction of sRALF and pRALF with FERONIA (Lan et al., 2023). In our models, we have also observed differential interactions and binding energies between stigmatic and pollen RALFs with FERONIA. Hence, our models complement the previously known experimental data (Lan et al., 2023). Despite these insights, our study could not fully decipher the competitive binding aspects between pollen and stigmatic RALFs. Further experiments, including mutation studies based on residue-level interaction details from our data, are required to elucidate these competitive dynamics comprehensively. Future advancements in high-resolution protein structure determination for these divergent ligand-receptor bindings hold promise for gaining deeper insights into the functional antagonism of these peptides.

Nevertheless, our study provides insights into the structural mechanisms of RALFs binding to FERONIA, which is a crucial step in understanding the molecular basis of hybridization in Brassicaceae. Additionally, this pipeline based on hybrid approach can be applied to investigate the binding of various flexible peptide-protein interactions using experimental restraints.

## Author contributions

HB: conceptualization, formal analysis, investigation, data curation, writing-original draft. KS: formal analysis, investigation, data curation. AS: formal analysis, investigation, writing-review and editing, supervision, funding acquisition. SS: conceptualization, investigation writing-review and editing, supervision, funding acquisition.

## Acknowledgments

We acknowledge the Ministry of Education for the Prime Minister Research Fellowship to HB and IITGN for fellowship to KS. We acknowledge support from DBT for the Ramalingaswami Re-entry fellowship and IITGN start-up grants to AS and SS.

## References

Berendsen, H. J. C., Postma, J. P. M., Van Gunsteren, W. F., DiNola, A., and Haak, J. R. (1984). Molecular dynamics with coupling to an external bath. The Journal of Chemical Physics 81:3684–3690.

Buchan, D. W. A., and Jones, D. T. (2019). The PSIPRED Protein Analysis Workbench: 20 years on. Nucleic Acids Res 47:W402–W407.

Dominguez, C., Boelens, R., and Bonvin, A. M. J. J. (2003). HADDOCK: A Protein-Protein Docking Approach Based on Biochemical or Biophysical Information. J. Am. Chem. Soc. 125:1731–1737.

Frederick, R. O., Haruta, M., Tonelli, M., Lee, W., Cornilescu, G., Cornilescu, C. C., Sussman, M. R., and Markley, J. L. (2019). Function and solution structure of the Arabidopsis thaliana RALF8 peptide. Protein Sci 28:1115–1126.

Haruta, M., Sabat, G., Stecker, K., Minkoff, B. B., and Sussman, M. R. (2014). A Peptide Hormone and Its Receptor Protein Kinase Regulate Plant Cell Expansion. Science 343:408–411.

Huang, P.-S., Ban, Y.-E. A., Richter, F., Andre, I., Vernon, R., Schief, W. R., and Baker, D. (2011). RosettaRemodel: A Generalized Framework for Flexible Backbone Protein Design. PLoS One 6:e24109.

Jones, D. T. (1999). Protein secondary structure prediction based on position-specific scoring matrices1. Journal of Molecular Biology 292:195–202.

Jumper, J., Evans, R., Pritzel, A., Green, T., Figurnov, M., Ronneberger, O., Tunyasuvunakool, K., Bates, R., Žídek, A., Potapenko, A., et al. (2021). Highly accurate protein structure prediction with AlphaFold. Nature 596:583–589.

Krissinel, E., and Henrick, K. (2005). Detection of Protein Assemblies in Crystals. In Computational Life Sciences (ed. R. Berthold, M.), Glen, R. C.), Diederichs, K.), Kohlbacher, O.), and Fischer, I.), pp. 163–174. Berlin, Heidelberg: Springer Berlin Heidelberg.

Kumar, S., Stecher, G., Li, M., Knyaz, C., and Tamura, K. (2018). MEGA X: Molecular Evolutionary Genetics Analysis across Computing Platforms. Mol Biol Evol 35:1547– 1549.

Lan, Z., Song, Z., Wang, Z., Li, L., Liu, Y., Zhi, S., Wang, R., Wang, J., Li, Q., Bleckmann, A., et al. (2023). Antagonistic RALF peptides control an intergeneric hybridization barrier on Brassicaceae stigmas. Cell 186:4773–4787.e12.

Letunic, I., and Bork, P. (2024). Interactive Tree of Life (iTOL) v6: recent updates to the phylogenetic tree display and annotation tool. Nucleic Acids Research Advance Access published April 13, 2024, doi:10.1093/nar/gkae268.

Li, S. C., and Ng, Y. K. (2010). Calibur: a tool for clustering large numbers of protein decoys. BMC Bioinformatics 11:25.

Liu, P., Haruta, M., Minkoff, B. B., and Sussman, M. R. (2018). Probing a Plant Plasma Membrane Receptor Kinase’s Three-Dimensional Structure Using Mass Spectrometry-Based Protein Footprinting. Biochemistry 57:5159–5168.

Liu, C., Shen, L., Xiao, Y., Vyshedsky, D., Peng, C., Sun, X., Liu, Z., Cheng, L., Zhang, H., Han, Z., et al. (2021). Pollen PCP-B peptides unlock a stigma peptide–receptor kinase gating mechanism for pollination. Science 372:171–175.

Parrinello, M., and Rahman, A. (1981). Polymorphic transitions in single crystals: A new molecular dynamics method. Journal of Applied Physics 52:7182–7190.

Pearce, G., Yamaguchi, Y., Munske, G., and Ryan, C. A. (2010). Structure-activity studies of RALF, Rapid Alkalinization Factor, reveal an essential--YISY--motif. Peptides 31:1973–1977.

Raveh, B., London, N., and Schueler-Furman, O. (2010). Sub-angstrom modeling of complexes between flexible peptides and globular proteins. Proteins 78:2029–2040.

Raveh, B., London, N., Zimmerman, L., and Schueler-Furman, O. (2011). Rosetta FlexPepDock ab-initio: Simultaneous Folding, Docking and Refinement of Peptides onto Their Receptors. PLOS ONE 6:e18934.

Sayers, E. W., Bolton, E. E., Brister, J. R., Canese, K., Chan, J., Comeau, D. C., Connor, R., Funk, K., Kelly, C., Kim, S., et al. (2021). Database resources of the National Center for Biotechnology Information. Nucleic Acids Res 50:D20–D26.

van Zundert, G. C. P., Rodrigues, J. P. G. L. M., Trellet, M., Schmitz, C., Kastritis, P. L., Karaca, E., Melquiond, A. S. J., van Dijk, M., de Vries, S. J., and Bonvin, A. M. J. J. (2016). The HADDOCK2.2 Web Server: User-Friendly Integrative Modeling of Biomolecular Complexes. J Mol Biol 428:720–725.

Vangone, A., and Bonvin, A. M. (2015). Contacts-based prediction of binding affinity in protein–protein complexes. eLife 4:e07454.

Waterhouse, A., Procter, J., Martin, D., Clamp, M., and Barton, G. (2009). Jalview version 2: A Multiple Sequence Alignment and Analysis Workbench. Bioinformatics (Oxford, England) 25:1189–91.

Xiao, Y., Stegmann, M., Han, Z., DeFalco, T. A., Parys, K., Xu, L., Belkhadir, Y., Zipfel, C., and Chai, J. (2019). Mechanisms of RALF peptide perception by a heterotypic receptor complex. Nature 572:270–274.

Xue, L. C., Rodrigues, J. P., Kastritis, P. L., Bonvin, A. M., and Vangone, A. (2016). PRODIGY: a web server for predicting the binding affinity of protein–protein complexes. Bioinformatics 32:3676–3678.

