## Supplemental information for "Structural insights into the recognition of RALF peptides by FERONIA receptor kinase during Brassicaceae Pollination"

**Methods**

**Phylogenetic analysis of various AtRALFs**

Sequences of 37 AtRALFs and *Nicotiana attenuata* RALF2 were derived from the GenBank NCBI database (see Table S1) (Sayers et al., 2021). Multiple sequence alignment of peptide sequences was created using the CLUSTAL algorithm with MEGA X software (Kumar et al., 2018). Sequence alignment was color-coded according to sequence conservation using the software Jalview (Waterhouse et al., 2009). Phylogenetic trees were constructed with the MEGA X software using the Maximum likelihood algorithm with default parameters. Bootstrapping was performed 1000 times. The tree was rooted using *Nicotiana attenuata* RALF2 as an outgroup. The inferred trees were visualized using iTOL (<https://itol.embl.de/>) (Letunic and Bork, 2024).

**Structure prediction of various AtRALFs**

The secondary structures of sRALF23/33 and pRALF11/26 were predicted using PSIPRED 4.0, available at the PSIPRED workbench (Jones, 1999; Buchan and Jones, 2019). Tertiary structures were predicted using the AlphaFold2 Colab server by providing the respective query sequences with pdb100 template mode. All other parameters were used as the default for tertiary structure prediction (Jumper et al., 2021).

**Molecular Docking of FERONIA-RALF Structures**

The experimentally derived structure for FERONIA was derived from the PDB database (PDB ID 6A5E) (Xiao et al., 2019). The models of RALF peptides predicted using AlphaFold2 colab were docked with the crystal structure of FERONIA using ambiguous and unambiguous restraints (see Table S2) in Haddock 2.4 online server. All other parameters were used as default for the initial docking (Dominguez et al., 2003; van Zundert et al., 2016). Missing loops in cluster representatives obtained from Haddock 2.4 were modeled and optimized using the Rosetta remodel algorithm using 1000 trajectories and “quick and dirty” flag; all other parameters were set to default (Huang et al., 2011).

**Refinement of FERONIA-RALF structures**

Initial models from Haddock output were optimized for side chain clashes and prepacked using the Rosetta pre-pack protocol with the default parameters and further refined (energy minimization) using the Rosetta pep refine protocol with and without low-resolution optimization (lowres_preoptimize flag). The number of models predicted was 500 (nstruct flag). All other parameters were used as default during refinement (Raveh et al., 2010).

**Flexible peptide docking of FERONIA-RALF complexes**

Refined models were renumbered, and fragment files (3mer and 9mer) were generated using the Rosetta Fragment online server. These inputs were used for flexible peptide docking using the FlexpepDock *ab initio* protocol guided by experimental restraints (see Table S2) (Raveh et al., 2010). Flags used were pep_refine and lowres_abinitio to predict 50000 models, which were clustered using Calibur (Li and Ng, 2010). Clustering was performed based on the root mean square deviation of RALFs in the models.

**Binding and interaction analysis**

The top 5 scoring models from the largest cluster were used for binding energy analysis using the Prodigy online server (Vangone and Bonvin, 2015; Xue et al., 2016). Molecular surface electrostatics was calculated for representative models using the Adaptive Poisson Boltzmann Solver (APBS) in PyMOL. The interactions between FERONIA and various RALFs were analyzed using the ePISA online server (Krissinel and Henrick, 2005).

**Molecular Dynamics Simulation**

All-atom molecular dynamics (MD) simulations were carried out for FERONIA-RALF11, FERONIA-RALF23, FERONIA-RALF26, and FERONIA-RALF33 systems using GROMACS version 2020.4. The Amber99sb-ildn force field was employed for these simulations, and the TIP3P water model was used to solvate the proteins in a dodecahedron box. Na+ and Cl- ions were added to maintain a concentration of 0.15 M.

The energy minimization was conducted using the steepest descent algorithm with 50,000 steps or a maximum force of 500 kJ/mol/nm on any atom. Following minimization, the system underwent equilibration in NVT ensemble for 1 ns, using a modified Berendsen thermostat to keep the temperature at 300 K (Berendsen et al., 1984). Subsequently, the system was equilibrated in a two-step process under the NPT ensemble: first for 1 ns using the Berendsen barostat (Berendsen et al., 1984) and then for another 1 ns using the Parrinello-Rahman barostat (Parrinello and Rahman, 1981), both at a pressure of 1 bar. Constant temperature of 300 K was maintained using modified Berendsen thermostat (V-rescale). Finally, a 20-nanosecond production run was performed for each of the four complexes individually in an NPT ensemble using the Parrinello-Rahman barostat with pressure at 1 bar and temperature at 300 K.

The root mean square deviation (RMSD), root mean square fluctuation (RMSF), and distance calculations were performed using in-built programs of GROMACS gmx rms, gmx rmsf, and gmx distance, respectively.

**Figure legends:**

**Supplementary Figure 1. Multiple sequence alignment of *A.thaliana* RALFs.**

(A) Multiple sequence alignment of AtRALFs and NaRALF mentioned in Table S1, color-coded based on sequence conservation. The darker the color, the more conserved the residue. Red box indicates conserved motifs.

(B) Multiple sequence alignment of stigmatic and pollen RALFs, color-coded based on sequence conservation. The darker the color, the more conserved the residue. Black box indicates residues that are conserved specifically in sRALFs (stigmatic RALFs), and red box indicates residues specifically conserved in pRALFs (pollen RALFs).

**Supplementary Figure 2. Secondary structure prediction of representative stigmatic and pollen RALFs**

PSIPRED predictions of secondary structures in pRALF11(A), sRALF23(B), pRALF26(C) and sRALF33(D). 3-State prediction is denoted as follows: C (coils), E (beta strands), and H (Alpha helix).

**Supplementary Figure 3. Tertiary structure prediction of representative stigmatic and pollen RALFs**

Tertiary structure predictions of pRALF11(A), sRALF23(B), pRALF26(C) and sRALF33(D) using AlphaFold2 Colab. Each structure is shown on the left, and their corresponding predicted aligned error (PAE) plots (middle) and predicted local distance difference test (pLDDT) plots are shown on the right. (Red: pRALF11, Green: sRALF23, Yellow: pRALF26, Orange: sRALF33)

**Supplementary Figure 4. Haddock cluster representatives of FERONIA-RALF complexes.**

Representative models from selected clusters of pRALF11(A), sRALF23(B), pRALF26(C), and sRALF33(D) in complex with FERONIA predicted by Haddock2.4. (Blue: FERONIA, Red: pRALF11, Green: sRALF23, Yellow: pRALF26, Orange: sRALF33)

**Supplementary Figure 5. Analysis of FERONIA-RALF complex models predicted using Rosetta FlexpepDock protocol.**

(A) Score versus root mean square deviation (RMSD) plot for top 5000 scoring models predicted using Rosetta FlexpepDock *ab initio* protocol.

(B) Score versus root mean square deviation (RMSD) plot for top 2 clusters of FERONIA-RALF complexes clustered using Calibur.

(C) Cluster representative models of various FERONIA-RALF complexes predicted using FlexpepDock *ab initio* and clustered using Calibur. (Blue: FERONIA, Red: pRALF11, Green: sRALF23, Yellow: pRALF26, Orange: sRALF33)

(D) Representative structures of various FERONIA-RALF models from the largest cluster, colored based on electrostatic potential as calculated by APBS. Red color indicates negative potential, and blue indicates positive potential. The darker the color, the more electrostatic potential.

**Supplementary Figure 6. Interaction between FERONIA and RALFs (stigmatic and pollen).**

(A) Structure of FERONIA-RALF23 showing polar contacts (left, shown as yellow dotted lines) and its magnified view(right). (Blue: FERONIA, Green: sRALF23)

(B) Structure of FERONIA-RALF33 showing polar contacts (left, shown as yellow dotted lines) and its magnified view(right). (Blue: FERONIA, Orange: sRALF33)

(C) Graph representing the distribution of charged residues across various stigmatic and pollen RALFs.

(D) Graph representing the distribution of proline across various stigmatic and pollen RALFs.

**Supplementary Figure 7. Flexibility and interaction analysis for FERONIA-RALF complex by MD simulations.**

(A)Root mean square fluctuation of C-α atoms in RALF-FERONIA complexes for 20 ns MD simulation. The residue on the left of the black dotted line belongs to FERONIA, and on the right belongs to RALFs.

(B)Distance between C-α atoms of D222 (FERONIA) and R30 (stigmatic sRALF23/33) along the trajectory.

(C)Distance between C-α atoms of N206 (FERONIA) and R30 (stigmatic sRALF23/33) along the trajectory.

**Supplementary Figure 8. Schematic workflow followed in this study for integrative modeling of FERONIA-RALF complex structure prediction.**

**Supplementary Figure 1:**

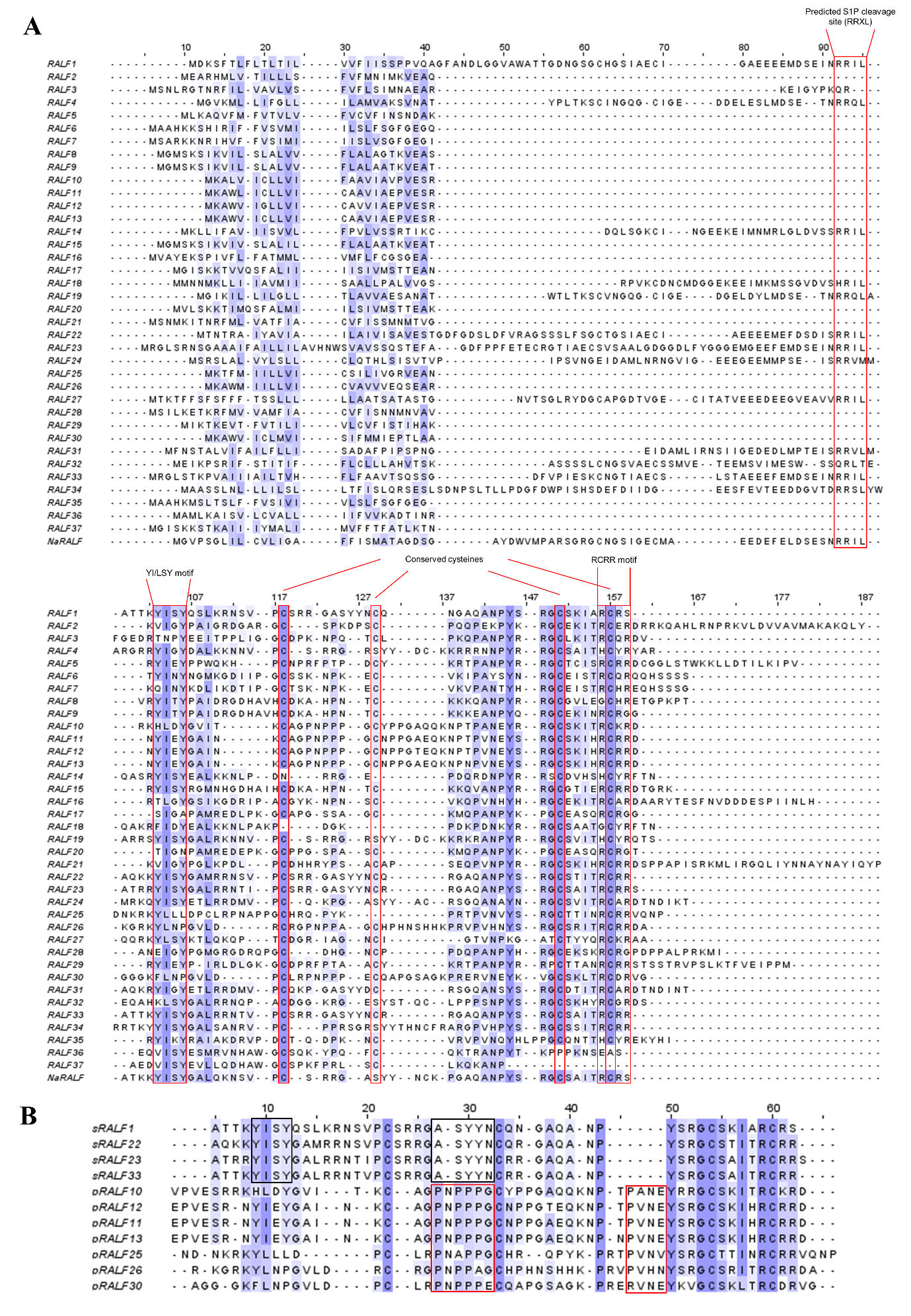

**Supplementary Figure 2:**

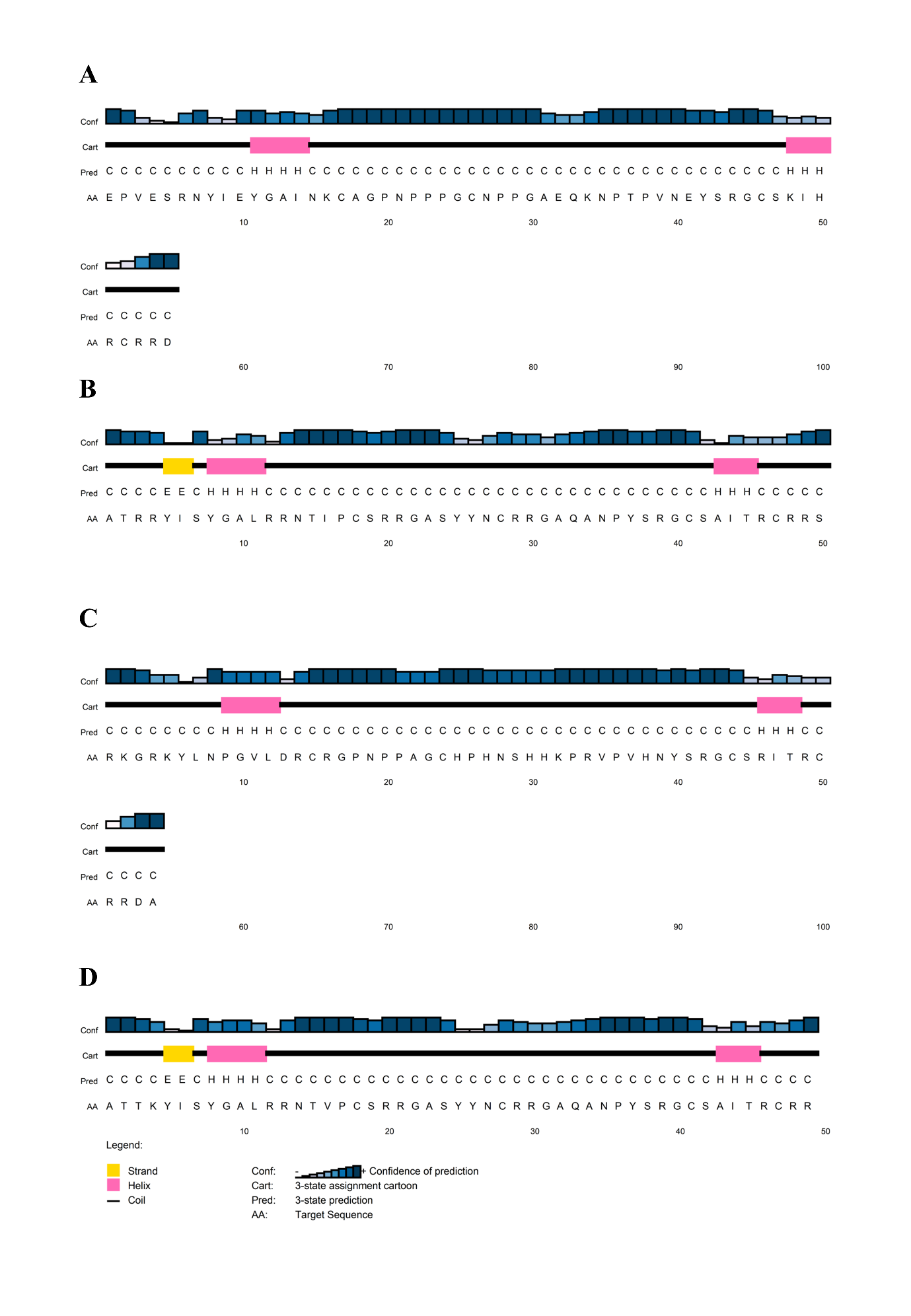

**Supplementary Figure 3:**

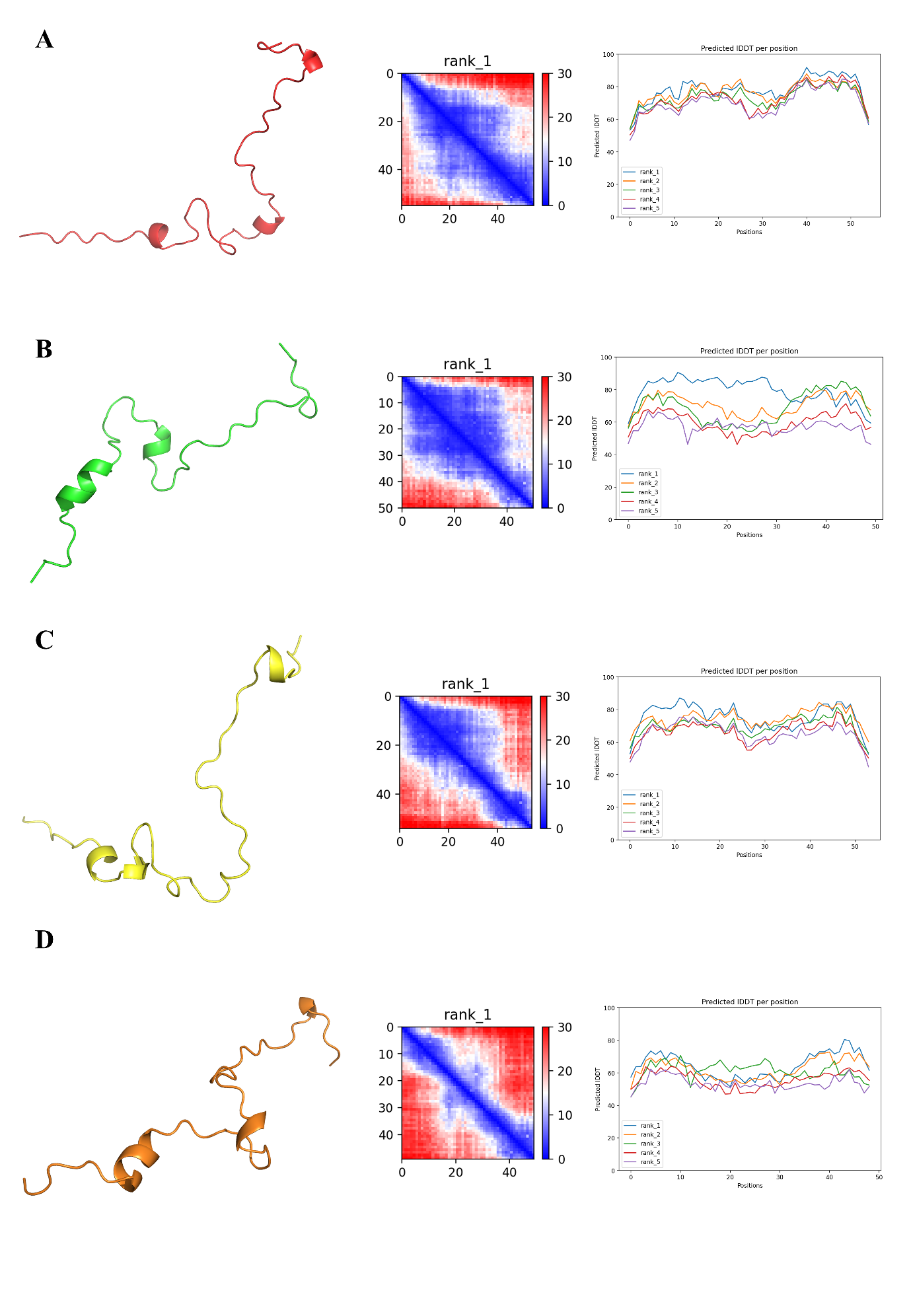

**Supplementary Figure 4:**

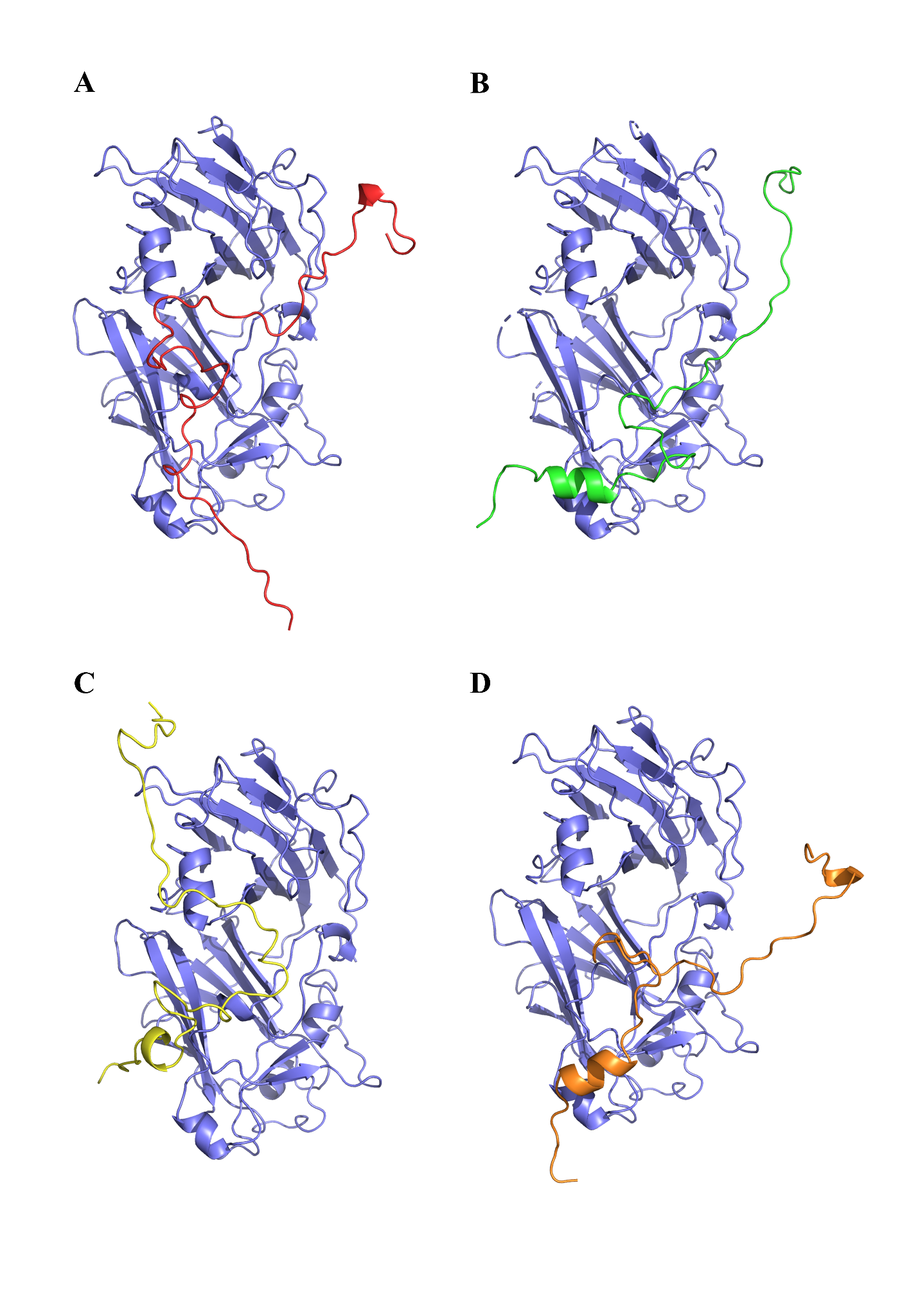

**Supplementary Figure 5:**

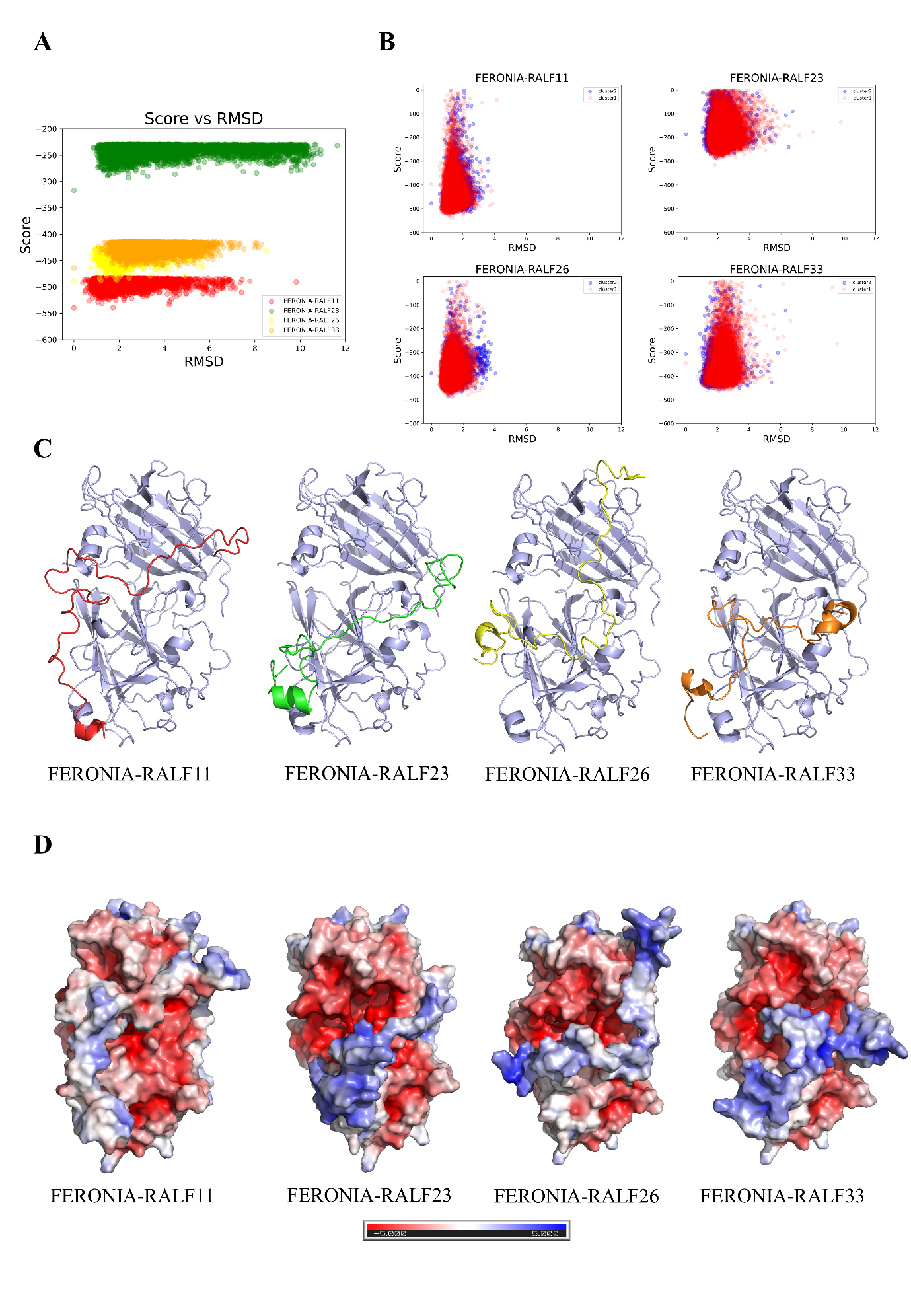

**Supplementary Figure 6:**

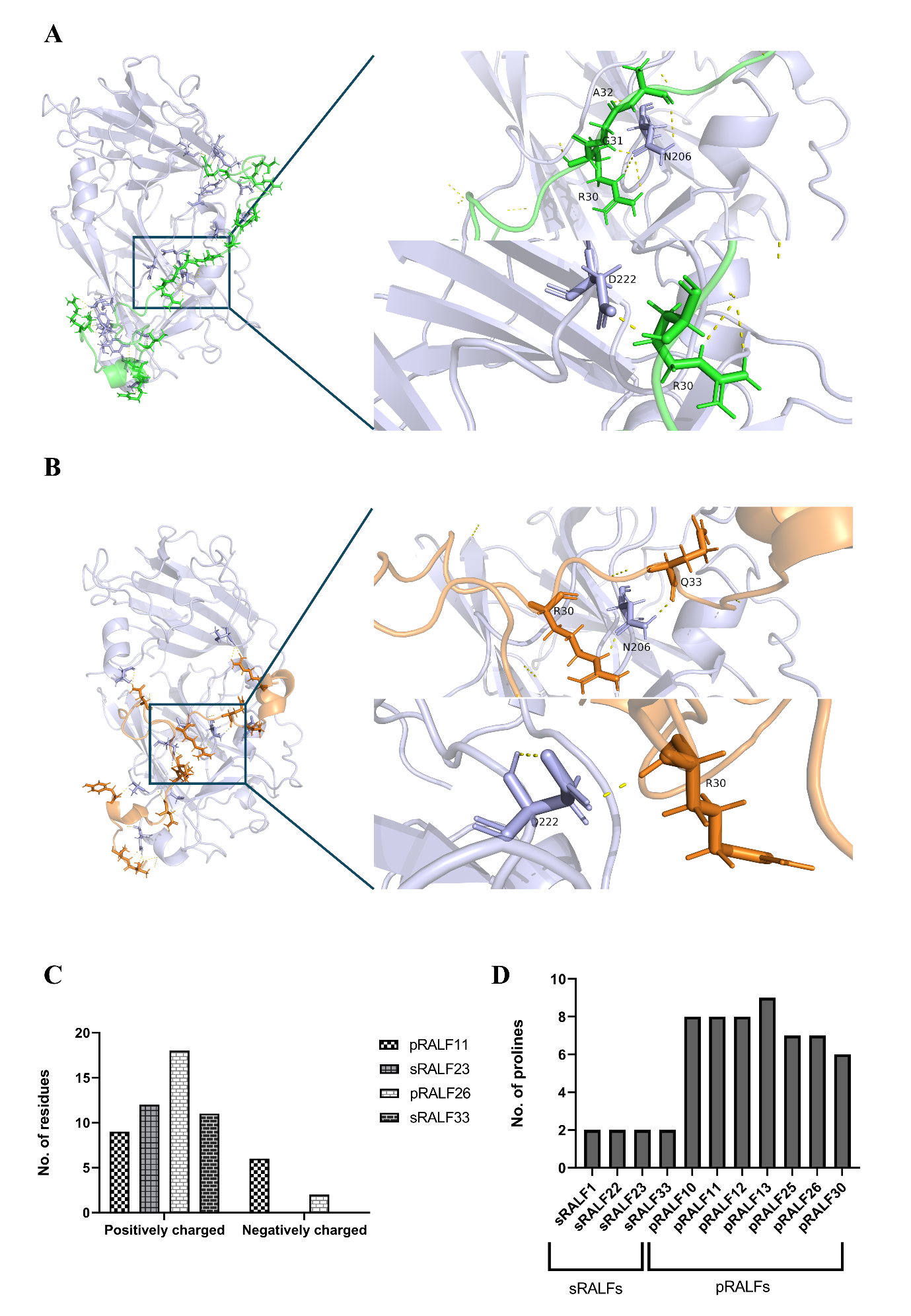

**Supplementary Figure 7:**

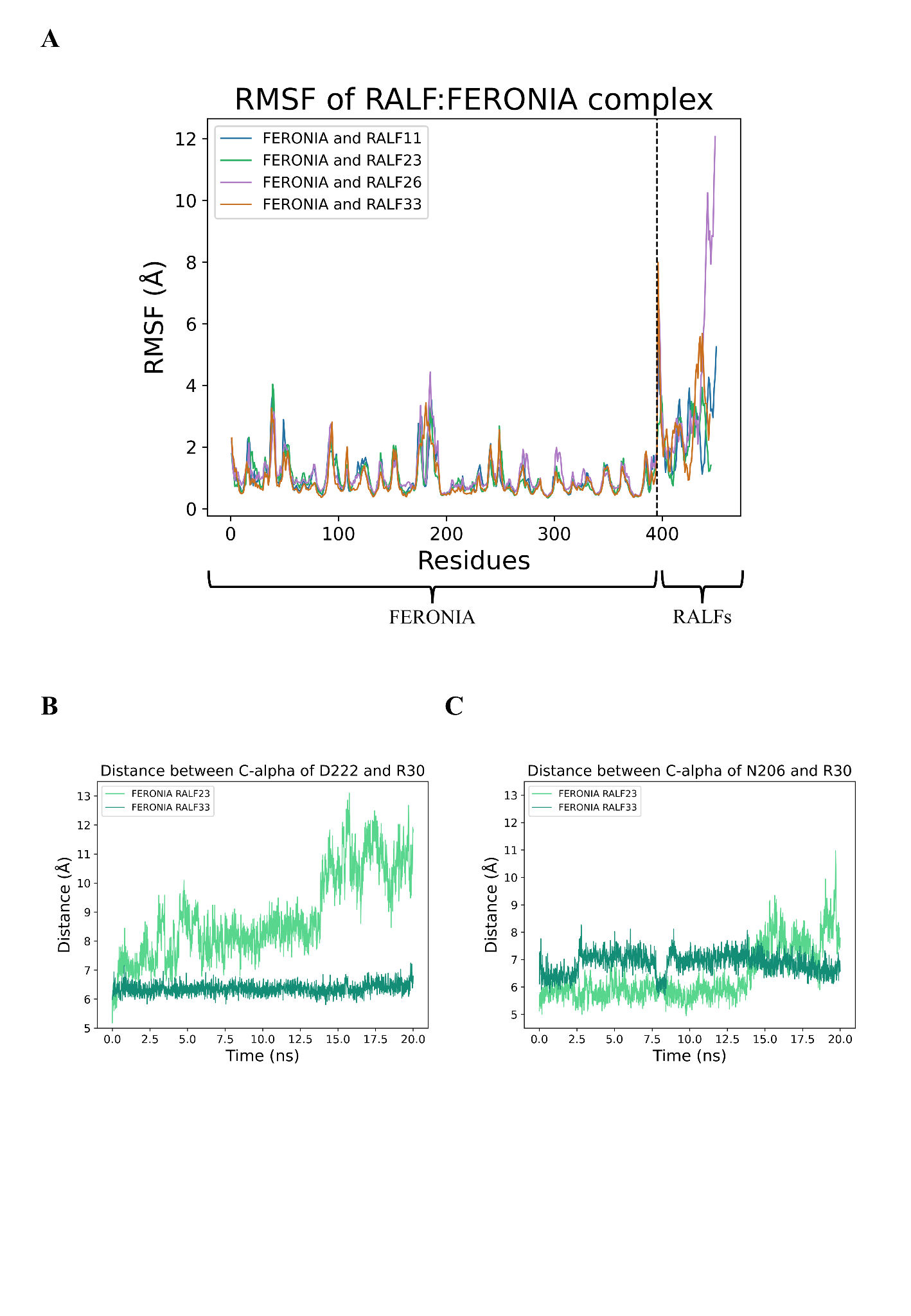

**Supplementary Figure 8:**

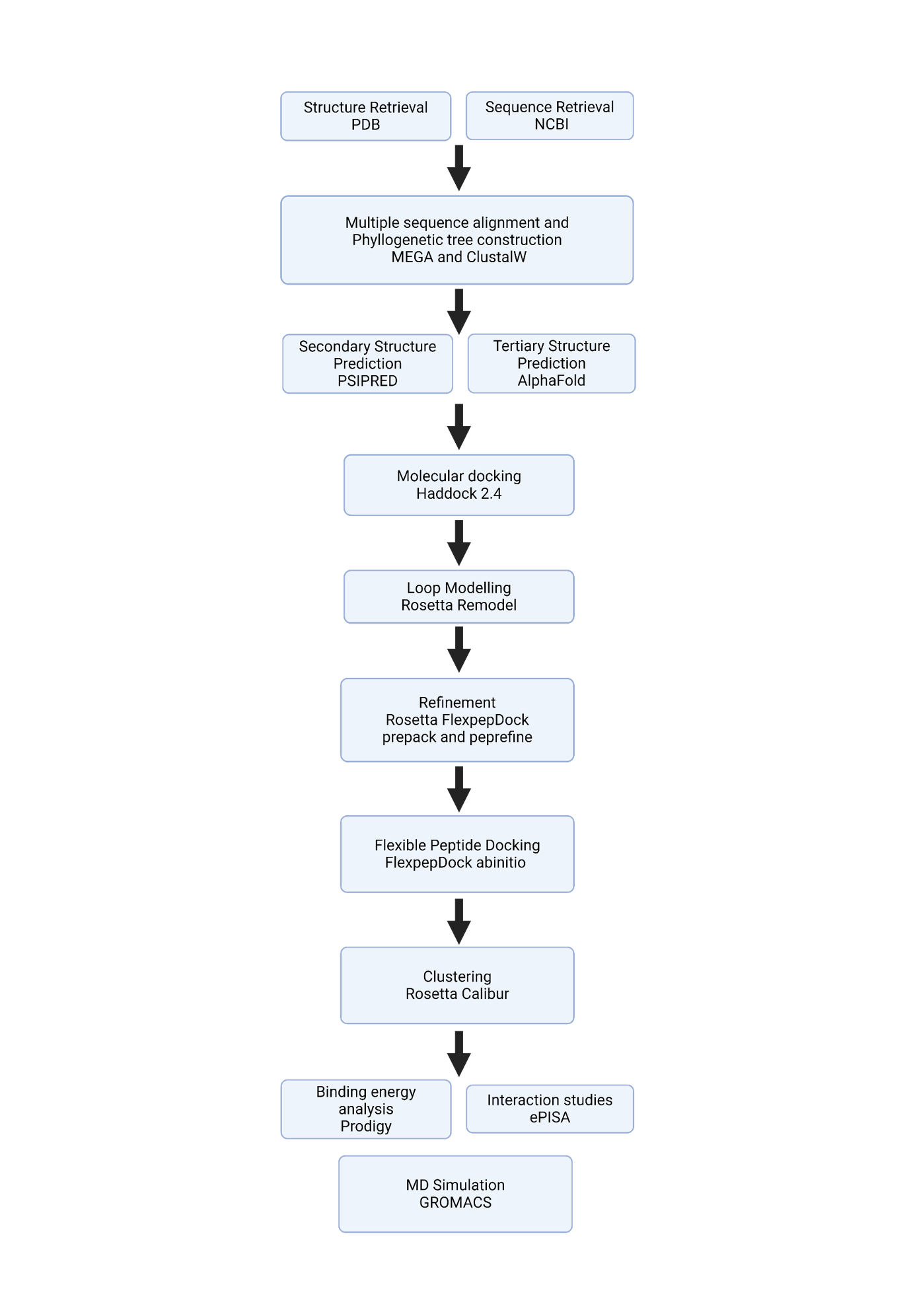

**Tables:**

**Table S1: Details of sequences used in this study.**

| Description | Nucleotide Accession | Uniprot Accession |
| --- | --- | --- |
| AtRALF1 | AT1G02900 | Q9SRY3 |
| AtRALF2 | AT1G23145 | A8MQ92 |
| AtRALF3 | AT1G23147 | A7REE5 |
| AtRALF4 | AT1G28270 | Q9FZA0 |
| AtRALF5 | AT1G35467 | A8MQI8 |
| AtRALF6 | AT1G60625 | A8MQM2 |
| AtRALF7 | AT1G60815 | A8MRD4 |
| AtRALF8 | AT1G61563 | Q1ECR9 |
| AtRALF9 | AT1G61566 | Q3ECL0 |
| AtRALF10 | AT2G19020 | O65919 |
| AtRALF11 | AT2G19030 | O64466 |
| AtRALF12 | AT2G19040 | F4ISE1 |
| AtRALF13 | AT2G19045 | F4ISE2 |
| AtRALF14 | AT2G20660 | Q9SIU6 |
| AtRALF15 | AT2G22055 | A8MQM7 |
| AtRALF16 | AT2G32835 | A8MRM1 |
| AtRALF17 | AT2G32885 | A8MR00 |
| AtRALF18 | AT2G33130 | O49320 |
| AtRALF19 | AT2G33775 | Q6NME6 |
| AtRALF20 | AT2G34825 | A8MQL7 |
| AtRALF21 | AT3G04735 | A8MRF9 |
| AtRALF22 | AT3G05490 | Q9MA62 |
| AtRALF23 | AT3G16570 | Q9LUS7 |
| AtRALF24 | AT3G23805 | Q9LK37 |
| AtRALF25 | AT3G25165 | Q9LSG0 |
| AtRALF26 | AT3G25170 | Q0V822 |
| AtRALF27 | AT3G29780 | Q9LH43 |
| AtRALF28 | AT4G11510 | Q9LDU1 |
| AtRALF29 | AT4G11653 | A8MQP2 |
| AtRALF30 | AT4G13075 | A7REH2 |
| AtRALF31 | AT4G13950 | Q2HIM9 |
| AtRALF32 | AT4G14010 | O23262 |
| AtRALF33 | AT4G15800 | Q8L9P8 |
| AtRALF34 | AT5G67070 | Q9FHA6 |
| AtRALF35 | AT1G60913 | A8MRK3 |
| AtRALF36 | AT2G32785 | A8MR74 |
| AtRALF37 | AT2G32788 | A8MRN0 |
| NaRALF | AY456269 | Q5QJ47 |

**Table S2: Restraints provided during molecular docking.**

*** represents ambiguity in the region of interaction**

| RALF | Restraint type | Residue number (FERONIA) | Residue number (RALF) | Source |
| --- | --- | --- | --- | --- |
| RALF11 | Unambiguous | 32 | 48 | (Liu et al., 2018) |
|  |  | 228 | 11 | (Xiao et al., 2019) |
|  |  | 229 | 11 |  |
|  |  | 230 | 11 |  |
|  |  | 230 | 14 |  |
|  |  | 229 | 15 |  |
|  |  | 230 | 15 |  |
|  |  | 232 | 15 |  |
|  | Ambiguous | 21 | * | (Liu et al., 2018) |
|  |  | 149 | * |  |
|  |  | 207 | * |  |
|  |  | 321 | * |  |
|  |  | 402 | * |  |
|  |  | * | 12 |  |
| RALF23 | Unambiguous | 32 | 43 | (Liu et al., 2018) |
|  |  | 228 | 8 | (Xiao et al., 2019) |
|  |  | 229 | 8 |  |
|  |  | 230 | 8 |  |
|  |  | 230 | 11 |  |
|  |  | 229 | 12 |  |
|  |  | 230 | 12 |  |
|  |  | 232 | 12 |  |
|  |  | 230 | 15 |  |
|  |  | 232 | 15 |  |
|  |  | 233 | 16 |  |
|  |  | 234 | 16 |  |
|  |  | 266 | 16 |  |
|  |  | 267 | 16 |  |
|  |  | 268 | 16 |  |
|  | Ambiguous | 21 | * | (Liu et al., 2018) |
|  |  | 149 | * |  |
|  |  | 207 | * |  |
|  |  | 321 | * |  |
|  |  | 402 | * |  |
| RALF26 | Unambiguous | 32 | 46 | (Liu et al., 2018) |
|  |  | 228 | 9 | (Xiao et al., 2019) |
|  |  | 229 | 9 |  |
|  |  | 230 | 9 |  |
|  |  | 230 | 12 |  |
|  |  | 229 | 13 |  |
|  |  | 230 | 13 |  |
|  |  | 232 | 13 |  |
|  | Ambiguous | 21 | * | (Liu et al., 2018) |
|  |  | 149 | * |  |
|  |  | 207 | * |  |
|  |  | 321 | * |  |
|  |  | 402 | * |  |
|  |  | * | 10 |  |
| RALF33 | Unambiguous | 32 | 43 | (Liu et al., 2018) |
|  |  | 228 | 8 | (Xiao et al., 2019) |
|  |  | 229 | 8 |  |
|  |  | 230 | 8 |  |
|  |  | 230 | 11 |  |
|  |  | 229 | 12 |  |
|  |  | 230 | 12 |  |
|  |  | 232 | 12 |  |
|  |  | 230 | 15 |  |
|  |  | 232 | 15 |  |
|  |  | 233 | 16 |  |
|  |  | 234 | 16 |  |
|  |  | 266 | 16 |  |
|  |  | 267 | 16 |  |
|  |  | 268 | 16 |  |
|  | Ambiguous | 21 | * | (Liu et al., 2018) |
|  |  | 149 | * |  |
|  |  | 207 | * |  |
|  |  | 321 | * |  |
|  |  | 402 | * |  |
|  |  | * | 9 |  |

**Table S3: Predicted FERONIA-RALF clusters by Haddock 2.4, their respective scores, and root mean square deviation (RMSD). Clusters highlighted in bold have been used for further analysis.**

| RALF | Cluster | Cluster size | Haddock score | RMSD  (Å) |
| --- | --- | --- | --- | --- |
| RALF11 | **1** | **171** | **-98.401** | **0.426** |
|  | 2 | 16 | -119.357 | 0.546 |
|  | 3 | 8 | -51.56 | 0.936 |
| RALF23 | **1** | **151** | **-107.772** | **1.653** |
|  | 2 | 12 | -42.443 | 2.519 |
|  | 3 | 8 | -88.282 | 1.932 |
|  | 4 | 4 | -48.924 | 2.027 |
|  | 5 | 4 | -121.426 | 1.073 |
| RALF26 | 1 | 39 | -62.843 | 0.573 |
|  | 2 | 37 | -57.706 | 1.35 |
|  | **3** | **25** | **-77.907** | **0.457** |
|  | 4 | 15 | -77.844 | 0.36 |
|  | 5 | 14 | -62.868 | 0.503 |
|  | 6 | 6 | -51.605 | 0.529 |
|  | 7 | 5 | -18.161 | 0.723 |
|  | 8 | 5 | -39.173 | 0.946 |
|  | 9 | 5 | -48.165 | 0.738 |
|  | 10 | 5 | -65.092 | 0.372 |
|  | 11 | 5 | -52.254 | 0.631 |
|  | 12 | 4 | -63.029 | 0.536 |
|  | 13 | 4 | -42.589 | 1.085 |
| RALF33 | **1** | **72** | **-76.495** | **0.575** |
|  | 2 | 29 | -51.366 | 0.502 |
|  | 3 | 27 | -73.292 | 0.464 |
|  | 4 | 14 | -43.884 | 0.285 |
|  | 5 | 14 | -46.707 | 0.499 |
|  | 6 | 8 | -26.871 | 1.875 |
|  | 7 | 7 | -37.983 | 0.653 |
|  | 8 | 7 | -58.016 | 0.704 |
|  | 9 | 6 | -40.847 | 0.846 |

**Table S4: Top 5 scoring models of FERONIA-RALF complexes predicted using FlexpepDock protocol.**

| Model No. | RALF | FlexpepDock Score | Average  score | RMSD with top-scoring model  (Å) | Average  RMSD  (Å) |
| --- | --- | --- | --- | --- | --- |
| 1 | RALF11 | -539.390 | -534.182 | 0.000 | 1.648 |
| 2 |  | -539.137 |  | 1.731 |  |
| 3 |  | -532.938 |  | 4.207 |  |
| 4 |  | -531.168 |  | 1.237 |  |
| 5 |  | -528.278 |  | 1.064 |  |
| 1 | RALF23 | -316.656 | -295.240 | 0.000 | 1.734 |
| 2 |  | -293.133 |  | 2.410 |  |
| 3 |  | -290.009 |  | 3.262 |  |
| 4 |  | -289.304 |  | 1.482 |  |
| 5 |  | -287.097 |  | 1.518 |  |
| 1 | RALF26 | -489.176 | -486.286 | 0.000 | 1.212 |
| 2 |  | -487.586 |  | 1.196 |  |
| 3 |  | -487.081 |  | 1.892 |  |
| 4 |  | -484.365 |  | 1.835 |  |
| 5 |  | -483.222 |  | 1.138 |  |
| 1 | RALF33 | -463.927 | -460.519 | 0.000 | 2.384 |
| 2 |  | -462.222 |  | 2.263 |  |
| 3 |  | -459.373 |  | 4.687 |  |
| 4 |  | -458.829 |  | 2.645 |  |
| 5 |  | -458.218 |  | 2.326 |  |

**Table S5: Size of clusters determined using Calibur.**

| Predicted structure | Cluster 1 size | Cluster 2 size |
| --- | --- | --- |
| FERONIA-RALF11 | 20042 | 3220 |
| FERONIA-RALF23 | 19924 | 4622 |
| FERONIA-RALF26 | 22515 | 2542 |
| FERONIA-RALF33 | 16182 | 2234 |

**Table S6: Binding energies predicted using Prodigy for the top 5 scoring FERONIA-RALF models of the largest cluster.**

| Model No. | FERONIA-RALF | FlexpepDock  Score | Average score | Binding energy  ΔG (kcal mol^-1^) | RMSD for RALFs  (Å) | Average  RMSD  (Å) |
| --- | --- | --- | --- | --- | --- | --- |
| 1 | FERONIA-Ralf11 | -539.390 | -532.812 | -13.9 | 0 | 0.938 |
| 2 |  | -539.137 |  | -12.6 | 1.731 |  |
| 3 |  | -531.168 |  | -12.1 | 1.237 |  |
| 4 |  | -528.278 |  | -14.6 | 1.064 |  |
| 5 |  | -526.089 |  | -12.9 | 0.660 |  |
| 1 | FERONIA-Ralf23 | -316.656 | -293.039 | -15.4 | 0 | 1.259 |
| 2 |  | -289.304 |  | -15.7 | 1.482 |  |
| 3 |  | -287.097 |  | -15.7 | 1.518 |  |
| 4 |  | -286.849 |  | -15.8 | 1.553 |  |
| 5 |  | -285.290 |  | -15.1 | 1.742 |  |
| 1 | FERONIA-Ralf26 | -489.176 | -484.884 | -11.8 | 0 | 1.243 |
| 2 |  | -487.586 |  | -11.3 | 1.196 |  |
| 3 |  | -484.365 |  | -11.6 | 1.835 |  |
| 4 |  | -483.557 |  | -11.1 | 1.138 |  |
| 5 |  | -479.735 |  | -12.9 | 2.047 |  |
| 1 | FERONIA-Ralf33 | -463.927 | -459.946 | -12.6 | 0 | 1.791 |
| 2 |  | -462.222 |  | -13.8 | 2.263 |  |
| 3 |  | -458.218 |  | -14.4 | 2.326 |  |
| 4 |  | -457.728 |  | -15.8 | 2.043 |  |
| 5 |  | -457.637 |  | -14.5 | 2.322 |  |

**Table S7. Polar contacts between various residues of representative FERONIA-RALF predicted models.**

| FERONIA | RALF11 | RALF23 | RALF26 | RALF33 |
| --- | --- | --- | --- | --- |
| D1 |  | Y37, S38 |  |  |
| S3 |  | R46 |  |  |
| E6 |  | R48 |  |  |
| K7 |  | T45, R46 |  |  |
| A16 |  |  | C44 |  |
| S17 |  |  | R42 |  |
| T20 | R44, G45 |  |  |  |
| D21 | Y42, G45 |  |  |  |
| T22 | Y42 |  |  | R48 |
| N24 | K48, D55 |  |  |  |
| K32 |  |  | A54 |  |
| E122 | Q33 |  |  |  |
| A123 | N27 |  |  |  |
| T125 | N35 |  |  | N27 |
| V164 |  | R48 |  |  |
| G205 |  |  | H27 |  |
| N206 |  | R30, G31, A32 |  | R30, Q33 |
| D207 |  | G31, Q33 |  |  |
| A212 |  | N35 |  |  |
| Y217 |  | R48 |  |  |
| D222 |  | R30 |  | R30 |
| Q224 |  |  | D13 | V16, P17 |
| P225 |  |  | R16 |  |
| F228 |  | R4 |  |  |
| A230 |  | R4 |  | Y8 |
| G233 |  | I16 |  | N14 |
| D258 |  |  | K32 | N35 |
| T269 |  | R13, N14 |  |  |
| N273 |  | N14 |  |  |
| N275 |  |  |  | T2, T3 |
| Y276 |  | N14 |  |  |
| N277 |  | A1 |  |  |
| T279 |  | A1 |  |  |
| N317 |  |  | K2 |  |
| D353 |  |  | R4 |  |
| Total no. of interactions | 8 | 16 | 9 | 11 |
